# Efficient inhibition of HIV using CRISPR/Cas13d nuclease system

**DOI:** 10.1101/2021.07.21.453273

**Authors:** Hoang Nguyen, Hannah Wilson, Sahana Jayakumar, Viraj Kulkarni, Smita Kulkarni

## Abstract

Recently discovered Clustered Regularly Interspaced Short Palindromic Repeats (CRISPR)/Cas13 proteins are programmable RNA-guided ribonucleases that target single-stranded RNA (ssRNA). CRISPR/Cas13 mediated RNA targeting has emerged as a powerful tool for detecting and eliminating RNA viruses. Here, we demonstrate the effectiveness of CRISPR/Cas13d to inhibit HIV-1 replication. We designed guide RNAs (gRNAs) targeting highly conserved regions of HIV-1. RfxCas13d (CasRx) in combination with HIV-specific gRNAs efficiently inhibited HIV-1 replication in cell line models. Furthermore, simultaneous targeting of four distinct sites in the HIV-1 transcript resulted in robust inhibition of HIV-1 replication. We also show the effective HIV-1 inhibition in primary CD4^+^ T-cells and suppression of HIV-1 reactivated from latently infected cells using the CRISPR/Cas13d system. Our study demonstrates the utility of the CRISPR/Cas13d nuclease system to target acute and latent HIV infection and provides an alternative treatment modality against HIV.

## Introduction

Globally 36.9 million people are infected with HIV. It is estimated that there are 5000 new infections every day (https://www.hiv.gov/hiv-basics/overview/data-and-trends/global-statistics). The development of combination therapy with multiple highly active antiretroviral drugs (HAART) has led to significantly more diverse treatment options and better quality of life for patients by mitigating against HIV-associated physical deterioration^1^. Efficacious HAART therapy reduces HIV-1 viral load to an undetectable level with a correspondingly high CD4^+^ T-cell count^2,3^. HAART therapy is a life-long commitment; disruption in therapeutic adherence can lead to the rapid development of resistant strains, complicating further treatment options and disseminating drug-resistant strains to new hosts^4^.

Furthermore, HAART has been shown to cause severe adverse events in many individuals, requiring empiric regimen changes for each, which may further affect adherence and, ultimately, the development of drug-resistant HIV-1 strains^5^. Attempts at HIV eradication have been mostly unsuccessful due to HIV genomic integration and maintenance as a latent reservoir in CD4^+^ T-cells ^6,7^ and macrophages^8,9^. The focus has been diverted towards a functional cure, facilitating viral inhibition without HAART intervention or eradicating the latent reservoir.

RNA interference (RNAi) represents one possible approach to achieve a functional cure for HIV. Small RNAs (si/shRNA) that use cellular RNAi machinery mediate HIV RNA decay efficiently. However, RNA structural constraints, HIV sequence diversity ^10-12^, viral protein-mediated suppression of cellular RNAi machinery^13-18^, and large-scale off-target effects ^19^ are some of the limitations of the RNAi approach. Therefore, we hypothesize that for HIV RNA-targeting strategy to be successful, it must have the following attributes: 1) The mechanism is independent of endogenous RNAi machinery. This significantly reduces the likelihood that the virus can inhibit the mechanism directly 2) Efficient and simultaneous targeting of multiple conserved sites. It has been predicted that the possibility of HIV-1 concurrently mutating at four or more conserved sites is negligible as it entails a great fitness cost to the virus^20-22^.

We used the recently discovered Clustered Regularly Interspaced Short Palindromic Repeats, CRISPR/RfxCas13d (CasRx) proteins to meet our above-hypothesized criteria. It is a programmable RNA-guided ribonuclease targeting single-stranded RNA (ssRNA) to target HIV-1 transcripts and genome. It has been reported that this ribonuclease is highly efficacious at degrading RNA in mammalian cells, with reduced dependency on RNA structure and significantly lower off-target effects than current siRNA/shRNA technologies^23-26^. We designed guide RNAs (gRNAs) targeting the conserved regions of HIV-1 group-specific antigen (gag), polymerase (pol), and central polypurine tract (cPPT) genes and showed that these gRNAs along with CasRx efficiently inhibit HIV replication. The inhibition of HIV replication was further improved when four distinct sites were targeted simultaneously. Our data show that CasRx protein combined with multiple gRNAs (poly-gRNA) strings simultaneously targeting four different conserved regions in HIV-1 degrades viral RNA efficiently in cell line models and primary cells and suppresses virus reemergence from latently infected cells. Our results suggest a potential utility of CasRx in controlling HIV-1 infection.

## Materials and Methods

### Plasmids

The CasRx, (pXR001; Addgene #109049), dCasRx (PXR002; Addgene #109050) plasmids, gRNA expression vectors (pXR003, pXR004; Addgene #109053 and #109054), Vesicular Stomatitis Virus glycoprotein (VSV-G) envelope expression vector (pMD2.G; Addgene #12259), lentiviral packaging plasmid (psPax2; Addgene # 12260), iRFP670 fluorescent reporter vector (piRFP670-N1; Addgene#79987) and pKLV2-U6gRNA5(BbsI)-PGKpuro2ABFP-W (Addgene #67974) were obtained from the Addgene repository (Watertown, MA, USA).

#### Construction of pBR43IeG-nef+-iRFP670 HIV vector

The open reading frame (ORF) sequence of iRFP670 amplified from piRFP670-N1 vector by polymerase chain reaction (PCR) using Q5® High-Fidelity DNA Polymerase (New England Biolabs, Ipswich, MA, USA) and primers encoding NcoI and XmaI digestion sites (Supplementary Table S1). The PCR product and pBR43IeG-nef+ (NIH AIDS reporter clone #11349) were then digested with NcoI and XmaI (New England Biolabs, Ipswich, MA, USA). The digested products were separated by 0.8% agarose gel electrophoresis and purified using the Monarch® DNA Gel Extraction Kit (New England Biolabs, Ipswich, MA, USA). The purified products were ligated using T4 DNA ligase (New England Biolabs, Ipswich, MA, USA). The ligated product was transformed into *Escherichia coli (E. Coli)* Stbl competent cells (New England Biolabs, Ipswich, MA, USA). The clones were picked and screened using colony PCR (Supplementary Table S1). The positive colonies were grown in Terrific Broth (Research Products International, Mount Prospect, IL, USA) containing 300 μg/mL ampicillin at 37 °C. The vector sequence was verified by Sanger sequencing (Genewiz, South Plainfield, NJ, USA).

#### Construction of pKLV2-U6-CasRx-(pre-gRNA)-PGKpuro2ABFP vector

The pKLV2-U6-CasRx-(pre-gRNA)-PGKpuro2ABFP vector was constructed using restriction enzyme digestion. The pre-gRNA cassette was PCR amplified using Q5® High-Fidelity DNA polymerase (New England Biolabs, Ipswich, MA, USA) to encode MluI and BamHI digestion sites as described in Supplementary Table S2. One microgram of this PCR product and pKLV-U6gRNA(BbsI)-PGKpuro2ABFP was digested with MluI and BamHI (New England Biolabs, Ipswich, MA, USA). The digested products were purified, ligated, transformed into *E. coli* Stbl competent cells (New England Biolabs, Ipswich, MA, USA), colonies were screened by PCR, positive clones were grown, and the vector sequence was further confirmed by Sanger sequencing as described above. Sanger Sequencing primers are described in Supplementary Table S1.

#### Construction of pVax-CasRx vector

CasRx-eGFP from pXR001 vector was PCR amplified and cloned into pVax1 (ThermoFisher Scientific, USA) by restriction enzyme cloning. The PCR product was digested with KpnI and XbaI (New England Biolabs, Ipswich, MA, USA) and ligated into pVaxI vector. The expression of CasRx-eGFP was evaluated by transient transfection in HEK293T derived Lenti-X™ cells (Takara Bio, Mountain View, CA, USA) cells, followed by visualization of eGFP by flow cytometry and confirmation of CasRx expression by qPCR (Primers listed in Supplementary Table S1).

#### Construction of CasRx-NES vector

Sequence from PXR001 vector was PCR amplified with specific oligonucleotide to exclude NLS sequence and add nuclear export signal of mitogen-activated protein kinase (MAPK). The rest of the backbone sequence of PXR001 was amplified in a separate PCR. Two DNA fragments with overlapping ends were prepared by consecutive PCR reactions with Q5 DNA polymerase. These fragments were then directly transformed into *E. coli* Stbl (New England Biolabs, Ipswich, MA, USA) for in vivo cloning as previously described^27^. Plasmid DNA extracted from resulting colonies was sequenced to confirm NLS signal removal and NES signal insertion in the CasRx-NES vector.

#### Single gRNA design and cloning

The single gRNAs were designed to target HIV transcript at highly conserved and siRNA targetable sites using the siVirus software (http://sivirus.rnai.jp/). These gRNAs were then confirmed for >70% genetic conservation with all HIV-1 variants within the Los Alamos National Laboratory HIV database (https://www.hiv.lanl.gov/). Those that match this criterion were cloned into the gRNA expression vector, PXR003 using the golden gate assembly. Primers used for gRNA are listed in Supplementary Table S2. 10 µM of each gRNA primer pair was phosphorylated and annealed in a 10 µL reaction containing 1 µL of 10x T4 DNA Ligase Buffer, 0.5 µL T4 polynucleotide kinase, and distilled water. Phosphorylation was carried at 37°C for 30 minutes. The primer pairs were annealed by increasing reaction temperature to 95°C for 5 minutes and allowing the reaction to cool down to 25 °C at a rate of 5°C/minute. The phosphorylated and annealed gRNAs were cloned into PXR003 using a Golder gate cloning reaction. Each reaction contained 1µM annealed crRNA guide oligonucleotides, 25 ng of PXR003, 0.5 µL BbsI (10U/ µL), 0.5 µL T4 DNA Ligase (400 U/ µL), 1 uL of 10x T4 DNA Ligase Buffer, and distilled water in total 10µl. The golden gate amplification was carried out in 30 repeating cycles of 37°C for 5 minutes and 23°C for 5 minutes. The golden gate mix was transformed into *E. coli* NEB Stbl competent cells (New England Biolabs, Ipswich, MA, USA) and sequenced with U6 promoter primer listed in Supplementary Table S2.

#### PolygRNA vectors

Four gRNAs, targeting four distant sites of HIV-1 genome or non-targeting (NT), along with direct repeat sequences, were collinearly arranged. The entire sequence was commercially synthesized (Genewiz, South Plainfield, NJ, USA) and cloned into pKLV-U6-CasRx-(pre-gRNA)-PGKpuro2ABFP vector using the golden gate assembly with some modifications. The modified 10 µL golden gate reaction contained 50 ng of synthesized polygRNA fragment, 25 ng of the pKLV-U6-CasRx-(pre-gRNA)-PGKpuro2ABFP, 0.5 µL BbsI (10U/µL), 0.5 µL T4 DNA Ligase (400 U/ µL), 1 µL of 10x T4 DNA Ligase Buffer, and distilled water.

### Cell culture

HEK293T derived Lenti-X™ cells (Takara Bio, Mountain View, CA, USA) and TZM-bl cells (ARP-8129; NIH AIDS Reagent Program) ^28^ were cultured in Dulbecco’s Modified Eagle Medium with high glucose, sodium pyruvate, and two mM L-Glutamine (Thermo Fisher Scientific) supplemented with 10% fetal bovine serum (ThermoFisher Scientific, USA) and antibiotic-antimycotic solution (ThermoFisher Scientific, USA). Cells were passaged to maintain <80% confluence. Jurkat Clone E6-1 (TIB-152™; ATCC®, Manassas, VA, USA) and HIV latency model J1.1 cells (ARP-1340; NIH AIDS Reagents Program) ^29^ were cultured in RPMI-1640 media supplemented with 2mM L-Glutamine, 10% bovine calf serum, and 1× antibiotic-antimycotic (Thermo Fisher Scientific). Primary CD4^+^T cells were isolated from peripheral blood lymphocytes (commercially available single-donor buffy coats from Innovative research https://www.innov-research.com/) and activated with Immunocult™ human CD3/CD28 T cell activator (Stemcell Technologies, Vancouver, Canada), in the RPMI-1640 media supplemented with 2mM L-Glutamine, 10% bovine calf serum, and 1× antibiotic-antimycotic (ThermoFisher Scientific, USA) and human recombinant IL-2 (Stemcell Technologies, Vancouver, Canada). Activated CD4+ T cells were transfected with pVax-CasRx and gRNA plasmids by nucleofection using Lonza 4d nucleofector. Eighteen hours after nucleofection, the cells were infected with HIV-iRFP virus particles. Virus replication was measured by flow cytometry after 48 hours of infection.

### Production of virus

The pBR43IeG-nef+-iRFP670 vector was transfected either by itself or with VSV-G envelop plasmid (PMD2.G) into Lenti-X™ cells using Transit-X2 (Mirus Bio, LLC) to make XR-4 tropic or VSV-G pseudotyped HIV-iRFP virus respectively. Antiviral supernatant was harvested 48 and 72 hours following transfection, centrifuged at 500g for 10 minutes at 4°C to remove cellular debris, and concentrated 10-fold using Lenti-X concentrator (Takara Bio, Mountain View, CA, USA). CasRx-GFP lentiviral particles were produced by co-transfecting plasmids encoding CasRx-GFP (PXR001; Addgene) with VSV-G envelope (pMD2.G; Addgene) and packaging (psPAX2; Addgene) into Lenti-X™ cells. Culture supernatants were harvested 48 hours post-transfection, clarified cell debris by centrifugation at 500g for 10 minutes at 4°C, and concentrated 10-fold using Lenti-X concentrator (Takara Bio, Mountain View, CA, USA). To obtain cells with stable CasRx-GFP expression, Lenti-X™ cells were transduced by spinoculation at 800g for 4 hours with CasRx-GFP lentivirus in growth media containing 8 µg/mL polybrene. Forty-eight hours after transduction, the top 5% cells expressing the highest levels of CasRx-GFP were sorted using BD FACS ARIA II (BD Bioscience, USA). The CasRx-GFP expressing cells were expanded and screened for CasRx expression using qPCR (primers listed in Supplementary Table S1).

#### CasRx-gRNA transfection and measurement of HIV replication by flow cytometry

Twenty-four hours before transfection, stable CasRx-GFP expressing Lenti-X™ cells (LRx) were plated at ∼80,000 cells per well in a 96 well flat bottom plate and incubated overnight to achieve ∼90% confluency before transfection. Two hundred nanograms (ng) of gRNA plasmids and 100 ng of pBR43IeG-iRFP670-nef+ plasmids were co-transfected per well. Each gRNA was transfected in three wells. Forty-eight hours after transfection, the cells were detached and made into a single cell suspension using Versene. HIV replication was measured using a BD Accuri C6 Flow Cytometer (BD Bioscience, USA) as percent HIV-iRFP670 expressing cells in HIV-guide RNA vs. the non-targeting control (NT) guide RNA transfected cells.

### Latency model

J1.1 cells were transfected with pVax-CasRx-GFP and gRNAs using Lonza 4d nucleofector. Eight hours post-nucleofection, the cells were stimulated with phorbol 12-myristate 13 acetate (PMA) and ionomycin (Cell stimulation cocktail, eBioscience, ThermoFisher Scientific, USA), and cell supernatants were collected after 24 hours.

### TZM-bl reporter assay

HIV production in the cell supernatants was measured using TZM-bl reporter cell line. Supernatants were added to the TZM-bl reporter cells, and the cells were incubated for 48 h. Cells were lysed, and luciferase activity was measured using Bright-Glo™ reagent (Promega, Madison, WI, USA) as previously described ^30^.

## Results

We employed the siVirus online tool (http://siVirus.RNAi.jp/)^31^ and the Los Alamos National Laboratory (LANL) database to select highly conserved sites in the HIV transcript. We selected twenty highly conserved target sites in HIV-1 transcript. The chosen sites are in the regions encoding gag, pol, protease (prot), integrase (int), cPPT, central termination sequence (CTS), and envelope (env). We designed gRNAs to target the selected sites in HIV (HIV-gRNAs; Figure 1A). We also designed a non-targeting control gRNA (NT) that does not target HIV-1 or any sequence in the human transcriptome (Supplementary Table S2). Alignment and comparison of gRNA sequences showed ≥70% conservation in the HIV-1 transcript sequences deposited in the LANL database (Supplementary Table S2).

**Figure 1.**
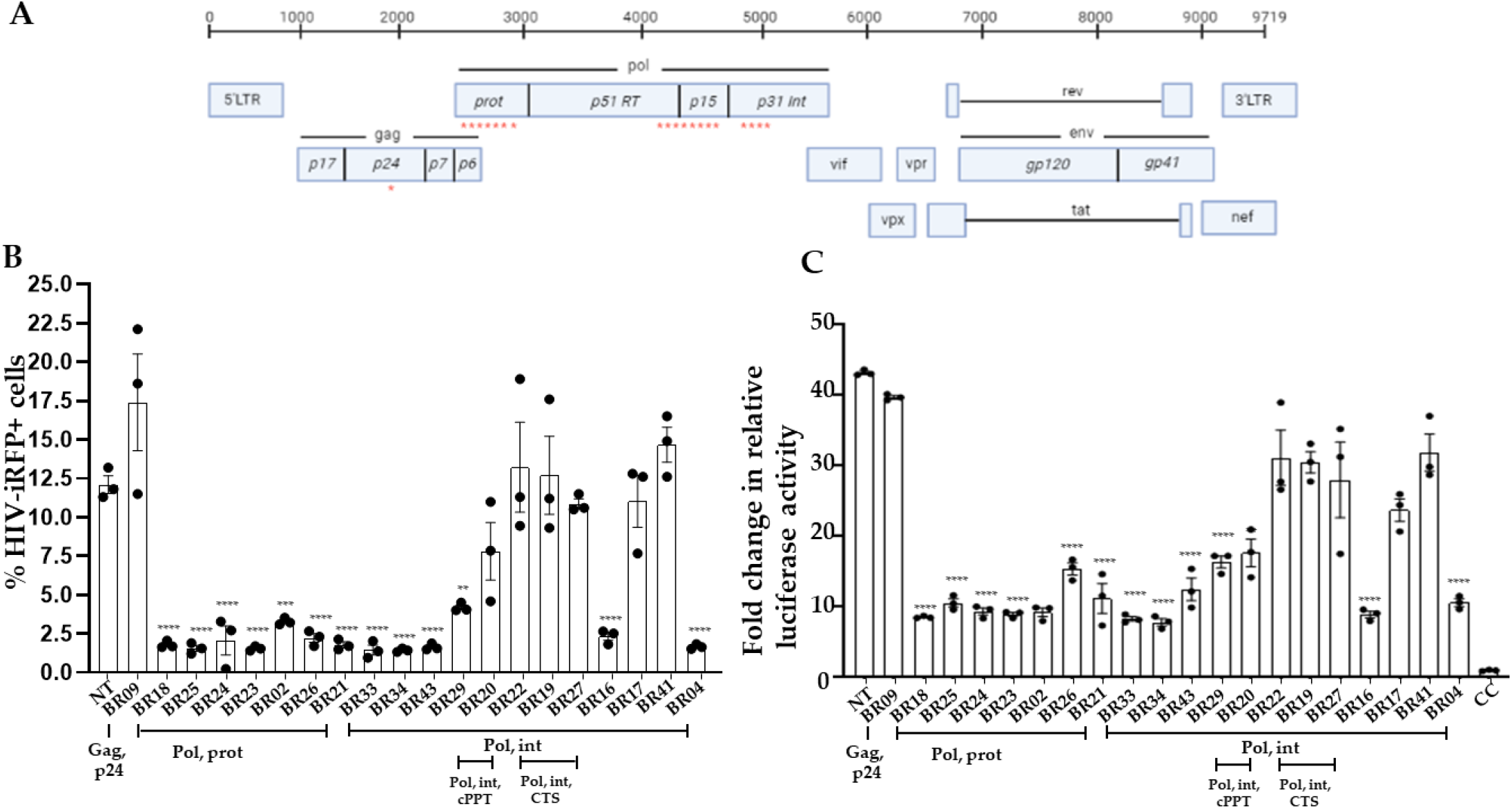
CRISPR/CasRx with HIV-specific single guide RNAs (gRNA) inhibits HIV replication. **(A)** Schematic representation of HIV genome and gRNA binding sites (red stars; HIV sequence compendium 2018) **(B)** Lenti-X™ cells with stable CasRx-GFP expression (LRx) cells were co-transfected with plasmid encoding gRNA and HIV-encoding iRFP670 fluorescent marker (HIV-iRFP). Forty-eight hours after transfection, expression of CasRx-GFP and HIV-iRFP were measured by flow cytometry. Positions of gRNA sequences in HIV transcript are indicated on the X-axis. **(C)** Supernatants collected from the LRx cells transfected with HIV-iRFP and HIV-gRNA or NT plasmids were added to the reporter cell line TZM-bl containing luciferase gene under HIV-LTR. Relative luciferase expression was measured after 48 h and presented as a fold change in relative luciferase activity normalized to cells only control (CC). The means ± s.e.m. are depicted as horizontal and vertical bars for each group, respectively. ANOVA tests with Bonferroni correction were used for statistical comparisons, and two-tailed *p* values are indicated. ****<0.0001; ***p<0.001, **p<0.01

To screen the HIV-targeting gRNAs, we developed an in vitro assay system comprising a stable cell line expressing RfxCas13d (CasRx-GFP) and a fluorescently labeled HIV-1 molecular clone. This allowed us to visualize the effect on HIV replication using flow cytometry. We used CasRx-GFP lentiviral construct encoding RfxCas13d fused to nuclear localization signal (NLS) at the N- and C-terminal and an enhanced green fluorescent protein (eGFP)^25^. A self-cleaving peptide sequence (2A) is inserted between Cas13d and eGFP coding sequences for simultaneous expression and cleavage of both proteins^25^. We transduced HEK-293 derived Lenti-X™ cell line with the NLS-CasRx-GFP lentiviral particles and isolated cells expressing the highest NLS-CasRx-GFP using a fluorescence-activated cell sorter (FACS; Supplementary Figure S1A). We confirmed CasRx expression in the sorted cells by quantitative PCR (qPCR; Supplementary Figure S1B). We refer to the Lenti-X™ cell line with stable expression of NLS-CasRx-GFP as LRx here.

We designed a molecular clone of HIV-1 expressing fluorescent tag to allow easy detection and quantification of virus infection, replication, spread, and inhibition by the CRISPR/CasRx system. We used dpBR43IeG-nef+ clone # 11349 (NIH AIDS reagent program)^32,33^, a full-length, chimeric, and replication-competent subtype B CXCR4-tropic vector (X4-tropic), derived from HIV-1 vector pNL4-3, designed to co-express eGFP and nef from a single bicistronic RNA. We replaced eGFP with a near-infrared fluorescent protein, iRFP670. We refer to this vector as HIV-iRFP (Supplementary Figure S2). HIV-iRFP vector can produce infectious HIV-1 particles that can infect T cells with CD4 and CXCR4 surface expression. The construct can also be used to generate pseudotyped lentiviral particles with the VSV-G envelope to infect a broader range of cell types.

We co-transfected the plasmids encoding NT or HIV-gRNAs, and HIV-iRFP vector in LRx cells, and 48 hours (h) post-transfection, measured viral replication by flow cytometry as percent iRFP expressing cells (Supplementary Figure S3). We also measured the production of infectious virions in the supernatant in HIV-gRNAs and NT transfected cells using a reporter cell line, TZM-bl. The TZM-bl cell line is a derivative of HeLa cells that express CD4, CCR5, and CXCR4 and contains an integrated reporter gene for firefly luciferase and *E. coli* β-galactosidase under the control of an HIV-1 long terminal repeat (LTR)^28^, permitting sensitive and accurate measurements of infection. Individual gRNAs targeting pol/prot (BR02, BR18, BR21, BR23, BR24, BR25, BR26), and pol/int (BR16, BR30, BR33, BR34, BR43, BR04, BR29) genes showed >70% reduction in HIV-1 replication as indicated by both the percent iRFP expressing cells (Figure 1B) as well as production of infectious HIV particles (Figure 1C) measured by TZM-bl reporter cell line compared to NT.

### Efficient inhibition of HIV using simultaneous targeting of four distant sites

Based on our preliminary analyses (Figure 1), we selected four gRNAs (BR04, BR23, BR34, BR29) targeting distinct regions in the HIV-1 transcript for poly-gRNA assembly (polyHIV). We transfected the LRx cells with plasmids encoding polyHIV or polyNT along with HIV-iRFP vector. The cells transfected with polyHIV showed >90% reduction in HIV-1 replication, as indicated by both the percent iRFP expressing cells (Figure 2A) and production of infectious HIV particles (Figure 2B) compared to polyNT control. We constructed a single lentiviral vector expressing CasRx and four guide RNAs. Lenti-X™ cell line was transduced with the CasRx-GFP-polyNT or CasRx-GFP-polyHIV lentiviral particles, and single cells expressing the highest GFP were isolated using FACS. The cells with stable expression of CasRx-polyNT (LRx-polyNT) or CasRx-polyHIV (LRx-polyHIV) were transfected with HIV-iRFP. The cells expressing CasRx-polyHIV showed a 99% reduction in HIV replication, as indicated by both the percent iRFP expressing cells (Figure 2C) and the production of infectious HIV particles (Figure 2D). The above experiments were repeated using catalytically inactive CasRx (dCasRx) and observed no change in HIV-RFP expression in HIV-gRNAs vs. NT gRNA transfected cells (Supplementary Figure S4). These data further confirmed efficient targeting and inhibition of HIV replication by CasRx and HIV-targeting gRNAs.

**Figure 2.**
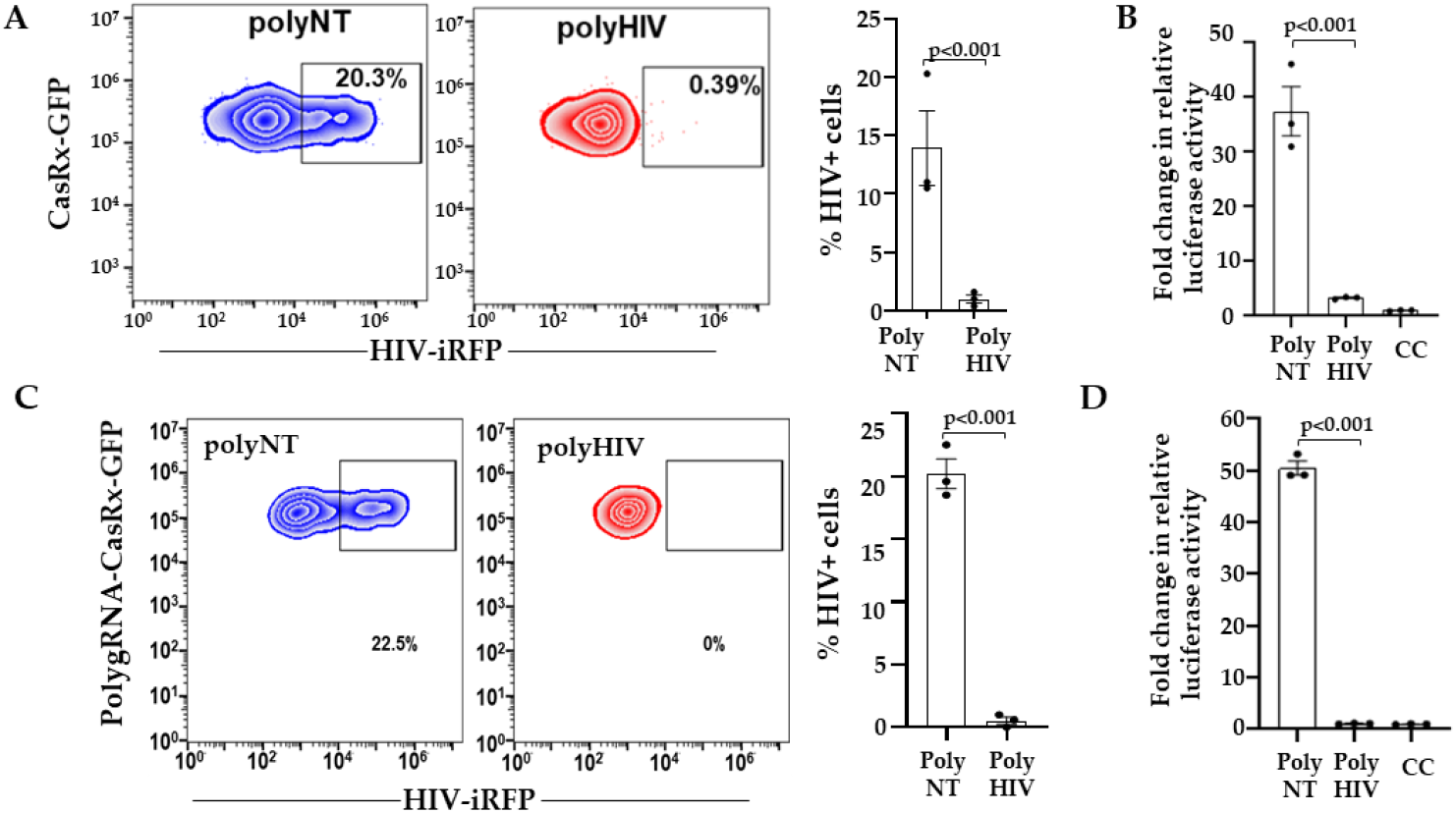
Inhibition of HIV RNA by simultaneous targeting of distinct regions. **(A)** LRx cells were transfected with plasmid poly-gRNAs targeting HIV (polyHIV; red) or non-targeting control (polyNT) and HIV-iRFP. Forty-eight hours after transfection, expression of CasRx-GFP and HIV-iRFP were measured by flow cytometry. **(B)** Supernatants collected from LRx cells transfected with HIV-iRFP and polyHIV or polyNT plasmids were added to the reporter cell line TZM-bl containing luciferase gene under HIV-LTR. Relative luciferase expression was measured after 48 h and presented as a fold change in relative luciferase activity normalized to cells only control (CC). **(C)** Lenti-X™ cells with stable expression of CasRx-polyNT (blue) or CasRx-polyHIV (red) were transfected with HIV-iRFP. Forty-eight hours after transfection, expression of CasRx-GFP and HIV-iRFP were measured by flow cytometry. **(D)** Supernatants collected from HIV-infected CasRx-polyHIV and CasRx-polyNT cells LRx were added to the reporter cell line TZM-bl containing luciferase gene under HIV-LTR. Relative luciferase expression was measured after 48 h and presented as a fold change in relative luciferase activity normalized to cells only control (CC). The means ± s.e.m. are depicted as horizontal and vertical bars for each group, respectively. The student’s t-test was used for statistical comparisons, and two-tailed *p* values are indicated.

To address if CasRx and gRNAs can target the incoming HIV in the cytoplasm, we engineered the CasRx protein to accumulate in the cytoplasm. We replaced the nuclear localization signals (NLS) with the MAPK nuclear export signal (NES) such that CasRx-NES protein is localized to the cytoplasm. We transfected Lenti-X™ cells with plasmids encoding CasRx-NES or CasRx-NLS and gRNAs. Twenty-four hours post-infection, the cells were infected with HIV-iRFP virus particles pseudotyped with Vesicular Stomatitis Virus G protein (VSV-G). Inhibition of virus infection in the polyHIV vs. polyNT transfected cells was assessed by flow cytometry 48 h post-infection. Cells transfected with nuclear (NLS) and cytoplasmic (NES) CasRx significantly reduced HIV replication in polyHIV gRNA transfected cells than polyNT transfected cells. However, the nuclear localization of CasRx was more effective in inhibiting HIV-1 replication than the cytoplasmic CasRx (Figure 3).

**Figure 3.**
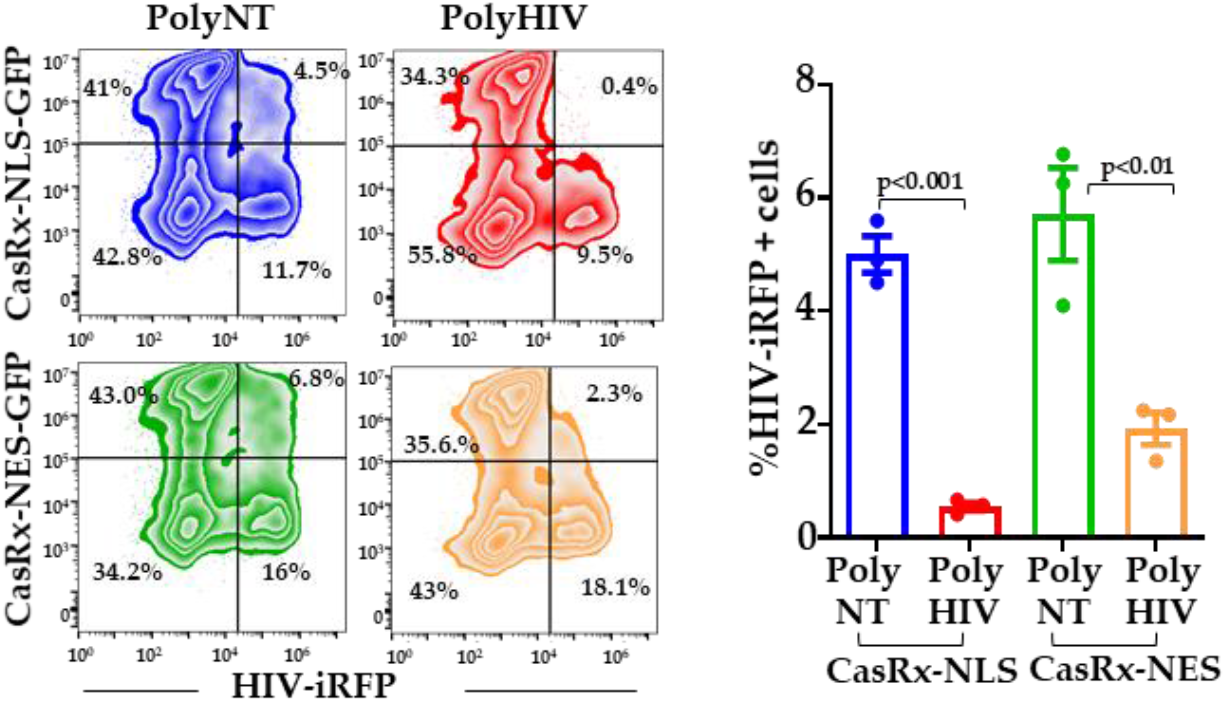
Nuclear localization of CasRx mediates optimal inhibition of HIV. Lenti-X^™^ cells were transfected with plasmid encoding CasRx-NLS-GFP, CasRx-NES-GFP, poly-gRNAs targeting HIV (polyHIV), or non-targeting control (polyNT). Twenty-four hours after transfections, the cells were infected with VSV-G pseudotyped HIV-iRFP. Forty-eight hours after infection, expression of CasRx-GFP and HIV-iRFP were measured by flow cytometry. The means ± s.e.m. are depicted as horizontal and vertical bars for each group, respectively. The student’s t-test was used for statistical comparisons, and two-tailed *p* values are indicated.

### CasRx inhibits HIV replication in primary CD4^+^ T cells

To facilitate better nucleofection efficiency in the primary CD4^+^ T cells, we cloned CasRx in a mammalian expression vector, pVax1 (pVax-CasRx). We transfected activated CD4^+^ T cells with plasmids encoding pVax-CasRx and gRNAs (polyHIV or polyNT) using nucleofection. The cells were infected with the X4-tropic HIV-iRFP virus particles 18 h post-transfection. HIV-iRFP expression was measured by flow cytometry 48 h after infection. The cells expressing pVax-CasRx and polyHIV significantly reduced HIV-iRFP expression than pVax-CasRx and polyNT (Figure 4).

**Figure 4.**
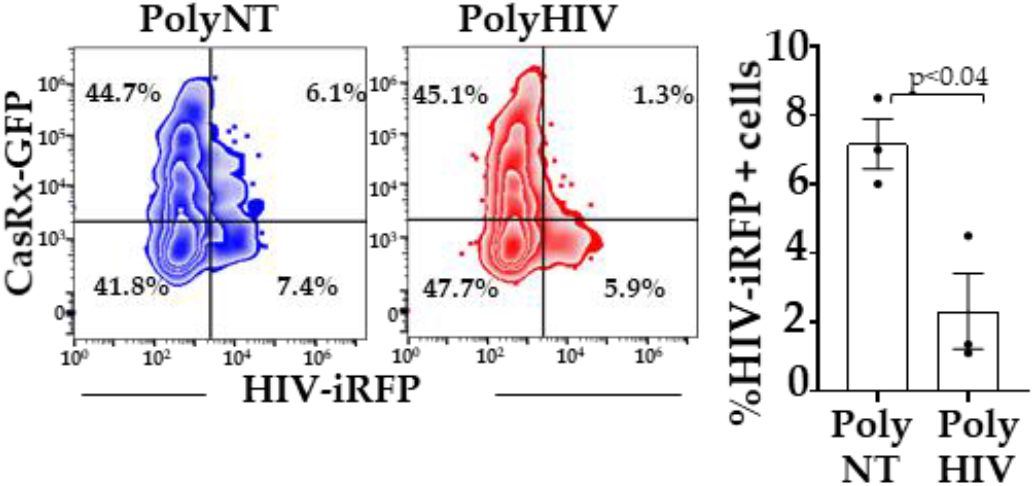
CasRx-silencing inhibited HIV replication in CD4^+^ T cells. We nucleofected pVax1-CasRx-GFP, non-targeting (polyNT), or HIV-targeting (polyHIV) gRNA plasmids in activated CD4^+^ T cells (n=3 distinct donors), infected the cells with HIV-iRFP, and measured HIV-iRFP expression 48 h post-infection by flow cytometry. The means ± s.e.m. are depicted as horizontal and vertical bars for each group, respectively. The paired t-test was used for statistical comparisons, and two-tailed *p* values are indicated.

### CasRx inhibits HIV expression from latent provirus

We used the human T cell line, J1.1 model, to test whether CasRx and HIV-gRNAs degrade HIV transcripts reactivated from HIV provirus. J1.1 is a chronically infected latent *cell* line cloned by limiting dilution from HIV-infected Jurkat cells and produces replication-competent HIV upon stimulation with latency reversal agents (LRA)^29^. We co-transfected J1.1 cells with plasmids encoding pVax-CasRx and gRNAs (polyHIV or polyNT) by nucleofection. The transfected cells were stimulated with PMA and ionomycin. We measured the production of infectious virions in the supernatant using a reporter cell line, TZM-bl (Figure 5A). Flow cytometry analyses showed that >80% of the nucleofected J1.1 cells expressed pVAX-CasRx (Figure 5B). We observed a significant reduction in HIV production in the cells transfected with polyHIV compared to polyNT (Figure 5C). These data indicate that CasRx-gRNAs can be used to suppress reactivated HIV from latently infected cells.

**Figure 5.**
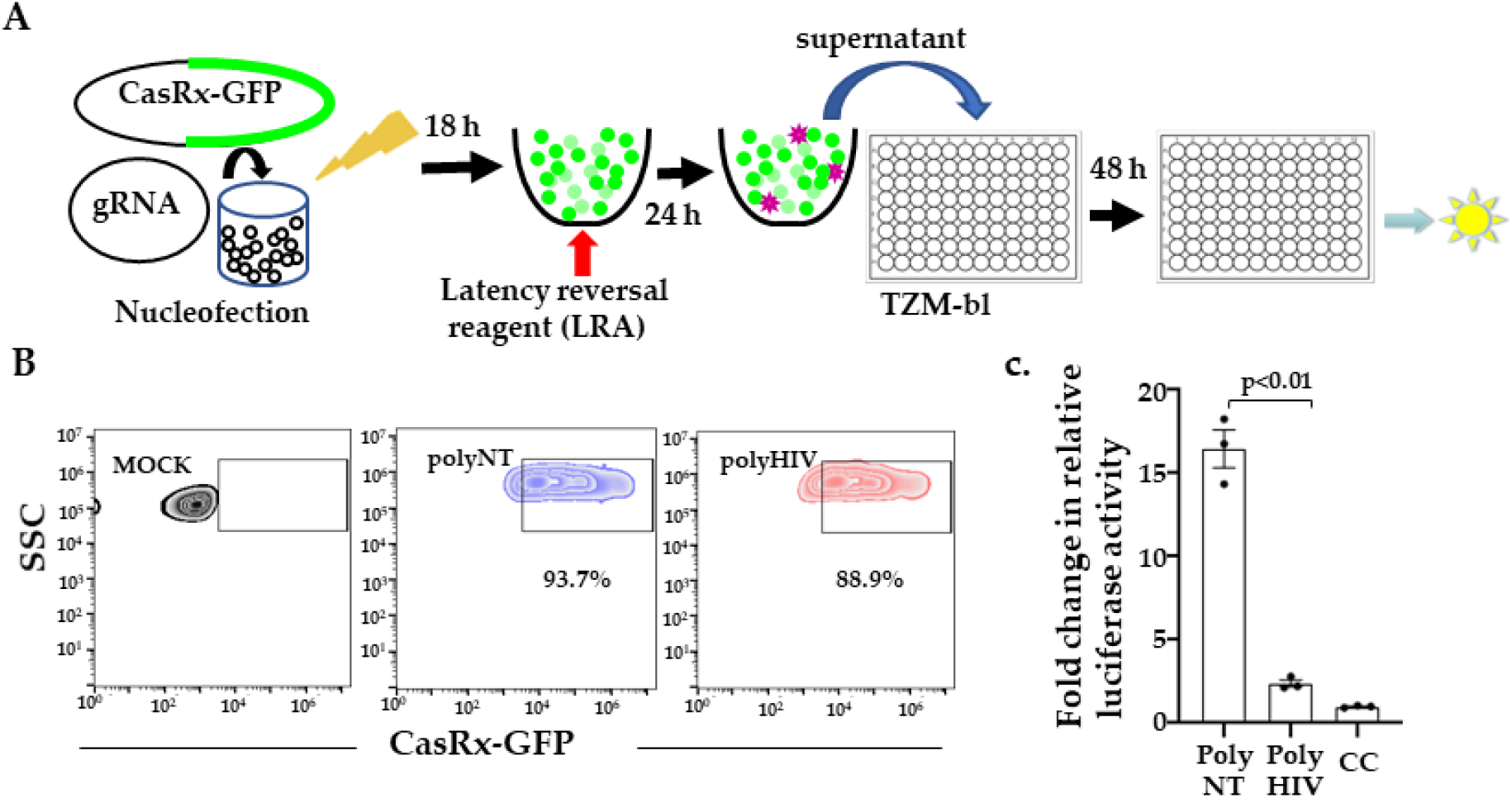
CasRx-silencing reduces HIV expression from latent provirus. **(A)** We nucleofected J1.1 cells with plasmids encoding gRNAs and CasRx-GFP and stimulated them with a latency reversal reagent (LRA; PMA), collecting the supernatants 24 h post-stimulation added to the TZM-bl cells. Relative luciferase expression was measured after 48 h and presented as a fold change in relative luciferase activity normalized to cells only control (CC). **(B)** Nucleofection of J1.1 cells with CasRx-GFP and gRNA plasmids. >80% of cells were transfected, indicated by GFP expression. **(C)** Supernatants from cells transfected with gRNAs targeting HIV (polyHIV) vs. non-targeting (polyNT) showed significantly reduced luciferase activity. The means ± s.e.m. are depicted as horizontal and vertical bars for each group, respectively. The student’s t-test was used for statistical comparisons, and two-tailed *p* values are indicated.

## Discussion

In this study, we present data demonstrating the use of CRISPR/CasRx nuclease system as an alternative treatment option against HIV. A major problem in developing effective countermeasures against HIV is viral diversity due to the error-prone nature of reverse transcriptase^34^. To overcome the diversity of HIV, several groups have developed vaccine candidates targeting the conserved regions^35-37^. This is because alterations in the conserved regions entail a fitness cost to the virus. We have used a novel RNA editing tool (CRISPR/CasRx) to target conserved regions of HIV and inhbit its replication in the current project. Out of the 20 gRNA screened, we identified 13 gRNAs highly effective in suppressing HIV replication. Most of the effective gRNAs were in the *pol* gene. HIV *pol* and *gag* are the two most conserved regions of the group M HIV clades^34^. Mutations in HIV *pol* gene in patients often result in reduced viral replicative fitness and pathogenecity^38,39^. To make our HIV-targetting approach more stringent, we generated a guide RNA construct (polyHIV, including BR04, BR23, BR29, BR34) that simultaneously target 4 distinct sites in highly conserved regions. With simultaneous targeting of four distinct conserved regions in HIV-1 sequence, we showed >90% inhibition of in vitro HIV replication. The gRNA target regions included the active site of the protease enzyme (BR23), the central polypurine tract (cPPT; BR29), the catalytic core domain of integrase (CCD, BR34), and the c-terminal domain (CTD; BR04) of integrase. Since these target sites are conserved in most of the circulating HIV-1 strains, we anticipate that this approach will be effective against all the HIV-1 clades.

The cellular location of CasRx nuclease is critical for efficient degradation of target RNA. It has been reported that cytoplasmic expression of CasRx is more effective against viruses that complete their life cycle in the cytoplasm. In our study, the nuclear localization of CasRx was found to be more effective against HIV than its cytoplasmic expression. Konerman et al. showed that nuclear localization signal increased CasRx activity in mammalian cells^25^. The ability to degrade RNA in the nucleus concurrently with transcription may confer CasRx superior gene knockdown efficiency. We believe this is advantageous over the siRNAs approach, as RNAi machinery is less effective in the nucleus.

The main target of HIV infection is the CD4^+^T cells^40^. We tested the efficacy of our antiviral cargo in primary CD4^+^T cells as well as lymphocytic cell lines. CasRx and gRNA plasmid nucleofection inhibited HIV replication in primary CD4^+^ T cells and suppressed the expression of viral RNA from activated latent HIV-1 provirus. These results indicate that CRISPR/CasRx system works in primary cells and can be effective in vivo. We are now developing methods to deliver our antiviral approach in animal models efficiently.

In vitro, RNAi against the virally transcribed RNA is highly effective in the short term at inhibiting HIV-1 replication ^20,41-45^. There are several limitations to the use of si/shRNAs. RNA structural constraints may limit the accessibility of the target sequence to si/shRNAs. HIV-1 escapes siRNA-mediated silencing by two mechanisms. HIV-1 is a highly mutagenic virus owing to the lack of proofreading activity within reverse transcriptase, RNA hypermutations induced by APOBEC3G, and viral genetic recombination ^10-12^. Thus sequence diversity of HIV limits the effective target sites to the highly conserved regions. Also, acquired mutations within or outside the target site alter its accessibility due to changes in sequence or RNA structure resulting in the escape of mutant virus^19^. In addition, HIV-1 infection has been shown to suppress endogenous RNAi machinery^13-18^. Furthermore, shRNA/siRNA therapies have the potential for large-scale off-target cleavage caused by mismatched-siRNA, induction of innate immune response caused by double-stranded RNA, overloading of host miRNA pathway^46,47^. The expression and activity of CRISPR/Cas13d are independent of cellular RNAi or microRNA machinery. Thus, it will remain effective in the HIV-infected cells with suppressed cellular RNAi pathways.

Type VI CRISPR nucleases have recently been discovered as site-specific RNA-guided, RNA-targeting effectors^23-26,48^. The nuclease activity of these proteins allows gene knockdown without genomic alteration. Cas13 proteins are divided into various subtypes, including Cas13a, Cas13b, and Cas13d. Cas13a and Cas13b require a protospacer flanking sequence (PFS), whereas Cas13d is PFS independent^25,49,50^. CRISPR/RfxCas13d (CasRx) is the optimized version of Cas13d by Konerman et al. ^25^. CasRx has superior knockdown efficiency compared to currently available methods, such as small hairpin RNA (shRNA) interference, dCas9-mediated transcriptional inhibition (CRISPRi), and Cas13a/Cas13b RNA knockdown^25^. Cas13a has been shown to reduce viral RNA of Dengue virus 2 (DENV2)^51^, HIV^52^, influenza A virus (FLUAV)^53^, and SARS-CoV2^53^. Cas13b has been used to target Chikungunya virus in a mosquito cell line^54^. Cas13d has been shown to inhibit SARS-CoV-2 infection in vitro and vivo^55^ and FLUAV in human lung epithelial cells^55^.

However, the off-target effects remain a major concern with RNA targeting strategies. In vitro, Cas13 orthologs have been demonstrated to have limited off-target effects compared to RNAi. However, recently Cas13a, used in most studies, showed collateral cleavage of untargeted cellular RNA in human glioma cells^56^. This collateral cleavage was not demonstrated for CasRx in mice and human neuronal lines, cementing it as a promising Cas13 ortholog for in vivo knockdown therapies^57^.

CRISPR/Cas9, a popular gene-editing tool, has been recently shown to treat HIV effectively in vitro ^58-67^ and animal models when administered in combination with antiretroviral therapy ^68,69^. This tool is being refined to cope with HIV genetic diversity, off-target effects, harmful effects of large and dramatic genome rearrangements, generation of escape variants due to Cas9-induced mutants ^62-65,70-72^. Combined CRISPR-Cas9 and RNAi mediated attack on distinct sequences of HIV-1 DNA and RNA showed additive inhibition of viral replication, delay, and even prevention of virus escape^73^. Given the limitations of RNAi, combination trial therapy with CRISPR/CasRx and Cas9 simultaneously targeting multiple distinct sequences in HIV RNA and DNA will effectively reduce the initial viral burden and inhibit viral escape. CasRx with multiple gRNAs targeting conserved sites in HIV could also be used as a combinatorial regimen with latency reversal agents in ‘shock and kill’ approaches to reduce the viral burden and de novo infection by the reactivated virus.

CRISPR/Cas13 system is entirely orthogonal to human cells (minimal potential interference with endogenous cellular processes). Cas13d/CasRx has an additional advantage of its small size that enables single vector Adeno-associated Virus (AAV) delivery of Cas13Rx and gRNA in vivo^25^. Recently, CasRx was functional when injected as RNA or ribonucleoprotein complexes with synthetic guide RNAs in zebrafish, medaka, killifish, mouse embryos, and drosophila^74^. AAV-mediated delivery of CasRx and gRNAs in mouse liver was successfully used to knock down target metabolic genes^75^. Further in vivo studies are warranted to assess the efficacy of this promising CRISPR/CasRx antiviral approach for a durable, and functional cure of HIV.

## Conclusions

CRISPR/CasRx nuclease system is a promising alternative tool against acute and chronic viral infections. We have demonstrated the effective use of this technology against HIV. Simultaneous delivery of a single vector with four gRNAs targeting distinct conserved regions of HIV is a robust approach to inhibit HIV replication. This approach will impede the generation of escape variants which is a significant problem in the field. This RNA-targeting strategy is easily programmable to target any host factor or RNA virus. We anticipate that it will be highly efficient against any RNA virus, especially highly mutable and infectious viruses. Our promising in vitro results warrant further optimization of in vivo delivery and assessment of efficacy, immunogenicity, and safety of this promising therapeutic modality against HIV.

## Author Contributions

Conceptualization, H.N., V.K., and S.K.; methodology, H.N., H.W., S.J., V.K., and S.K.; writing—original draft preparation, H.N.; writing—review and editing, H.N, H.W. S.J. V.K, and S.K.; supervision, S.K.; project administration, S.K.; funding acquisition, V.K., and S.K. All authors have read and agreed to the submitted version of the manuscript.

## Acknowledgments

We thank Dr. Vida Hodara and the Texas Biomed Biology core for cell sorting.

---

**Table S1.**
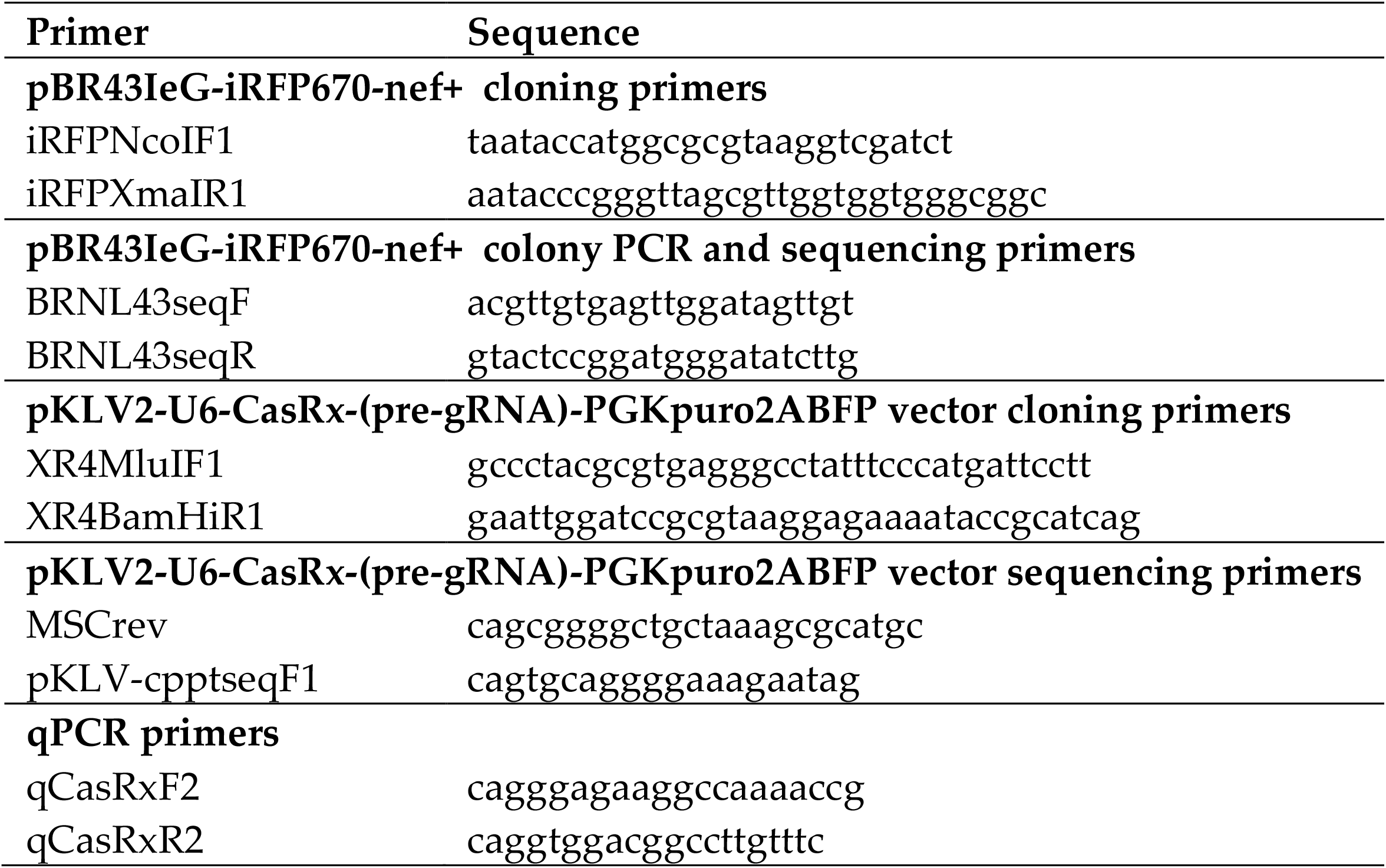
Oligonucleotide primers.

**Table S2.**
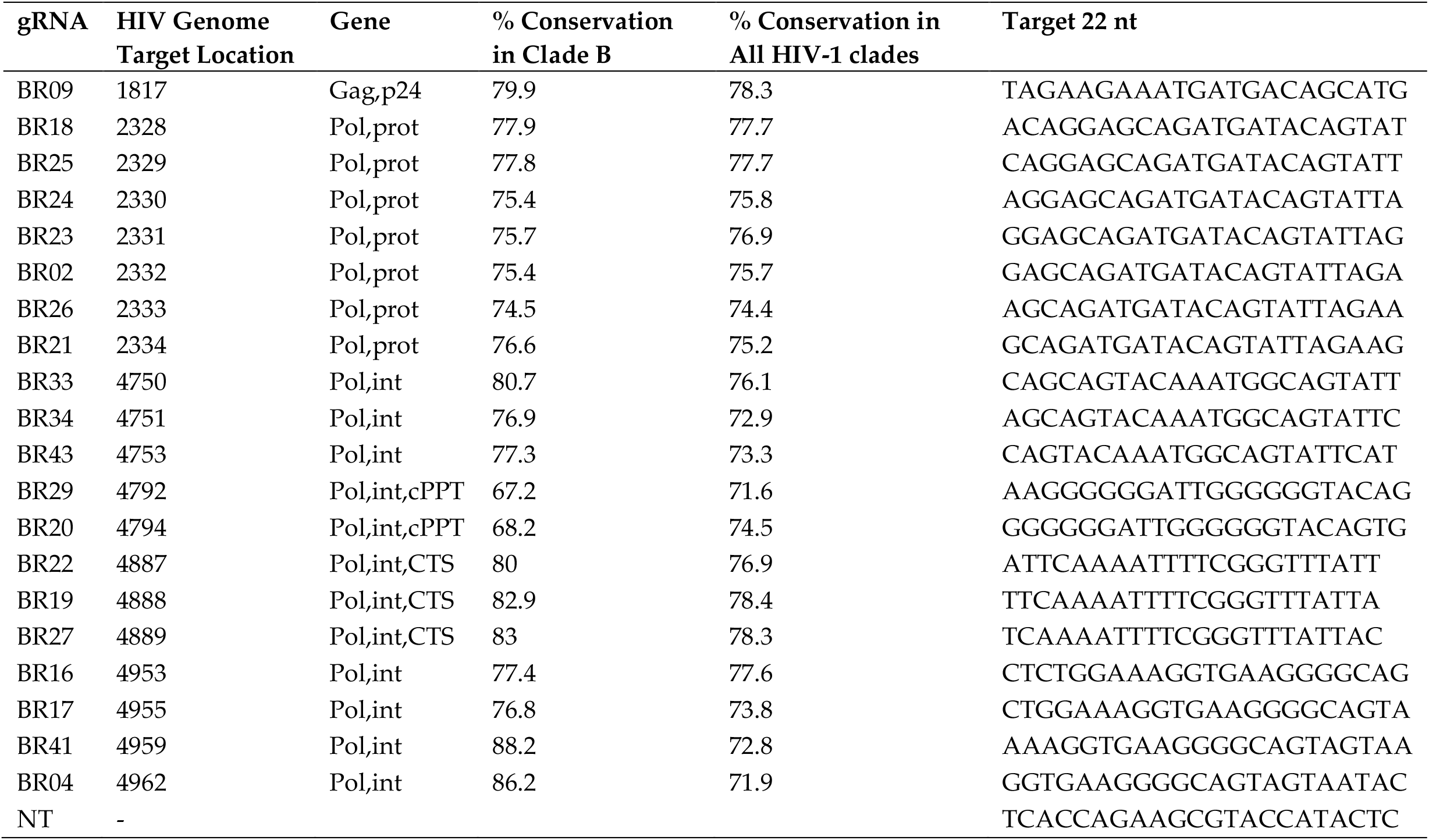
guide RNA design.

**Supplementary Figure S1.**
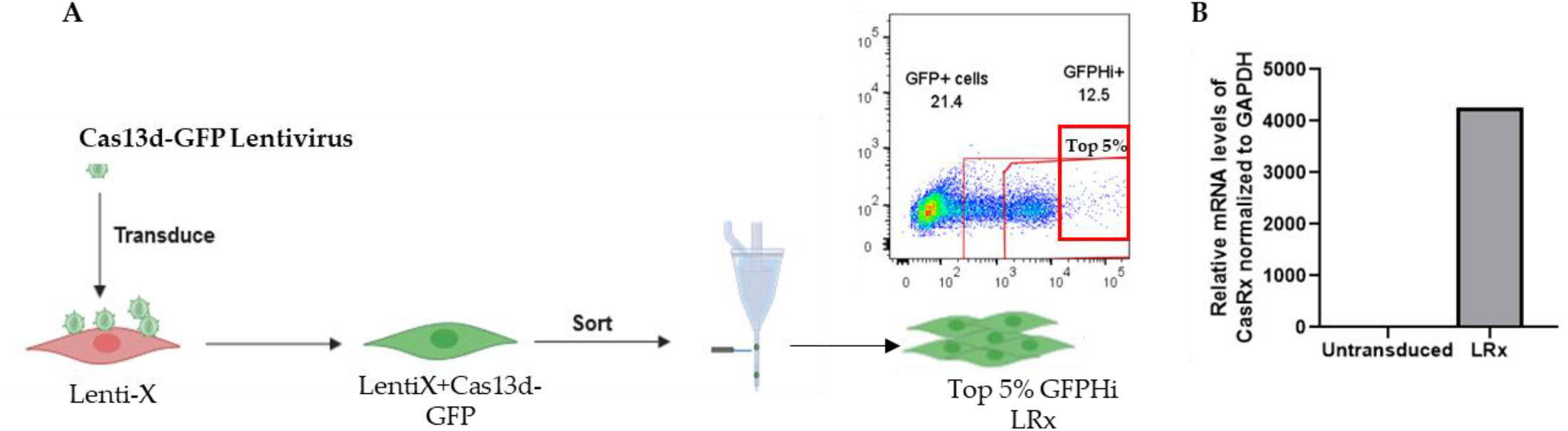
(A) Gaiting strategy. Lenti-X™ cell line was transduced with Cas13d-GFP lentiviral particle. Over twenty one percent cells expressed GFP (GFP+), 12.5% cells expressed high levels of GFP (GFPHi+). Cells expressing highest levels of GFP Cells expressing highest levels of GFP (Top5%) were sorted by FACS and further expanded in vitro culture. These cells with stable expression of Cas13d-GFP are termed LRx cells. (**B)** Expression of CasRx in LRx cells is confirmed by quantitative PCR (qPCR).

**Supplementary Figure S2.**
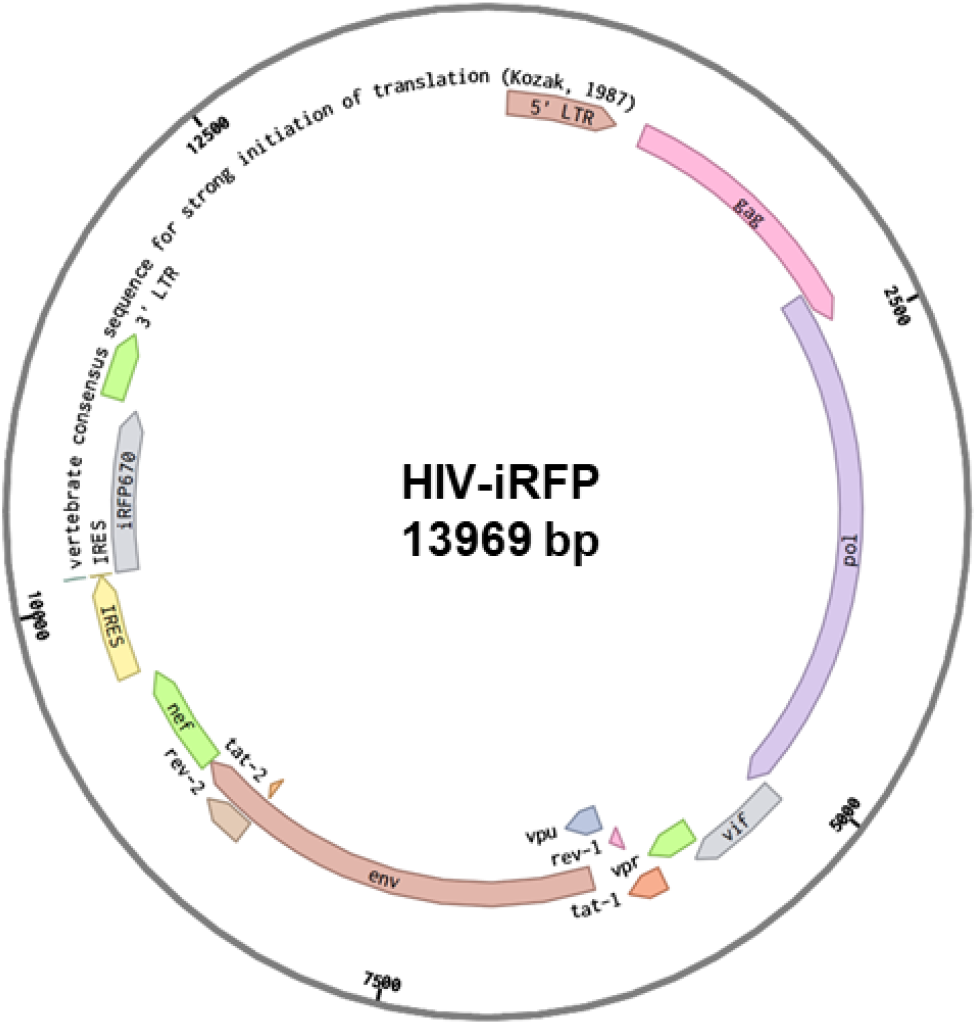
Plasmid design for expression of HIV-1 encoding fluorescent tag. A full-length, chimeric and replication competent subtype B, CXCR4-tropic HIV-1 vector (dpBR43IeG-nef+ clone # 11349; NIH-AIDS reagent, which co-express eGFP and nef from a single bicistronic RNA was modified to replace eGFP by a near infra-red fluorescent protein iRFP670. This vector henceforth referred to as HIV-iRFP.

**Supplementary Figure S3.**
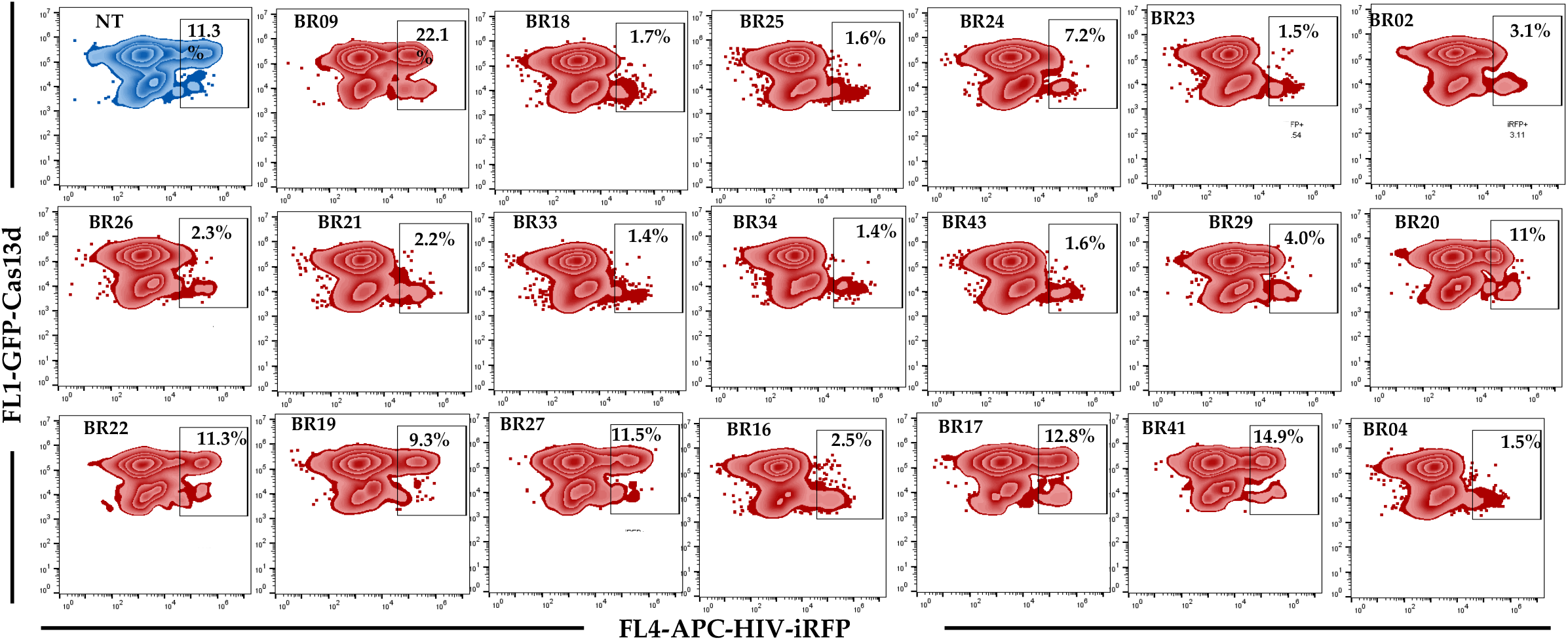
Measurement of viral replication by flow cytometry. Lenti-X™ cells with stable expression of CasRx-GFP (LRx) were transfected with plasmids encoding molecular clone of HIV co-expressing iRPF-670 fluorescent tag (HIV-iRFP) and HIV-specific guide RNAs (HIV-gRNA, red) or a non-targeting control (NT, blue). Forty eight hours after infection, expression of CasRx-GFP and HIV-iRFP were measured by flow cytometry. The cells transfected with HIV-targeting gRNA (red) inhibited HIV as compared to the cells transfected with Non-targeting gRNA (NT, blue).

**Supplementary Figure S4.**
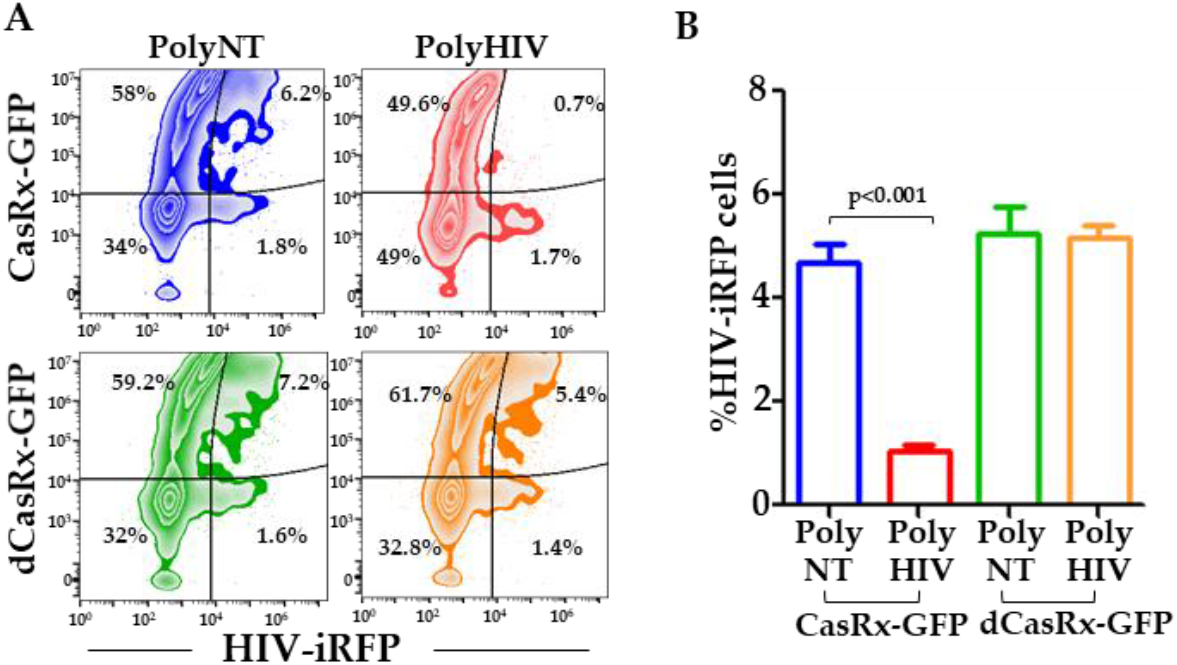
Catalytically inactive CasRx (dCasRx) with HIV-specific guide RNAs does not affect HIV replication. (**A**) Lenti-X ™ cells were co-transfected with plasmid encoding CasRx-GFP or dCasRx-GFP, guide RNAs (gRNA) and HIV-iRFP. Forty eight hours after infection, expression of CasRx-GFP/dCasRx-GFP and HIV-iRFP were measured by flow cytometry. (**B)** In the cells expressing CasRx-GFP and polyHIV (red) the percentage of HIV-iRFP+ cells were significantly reduced compared to CasRx-GFP and polyNT (blue). However, percentage of HIV-iRFP cells were similar in dCasRx-GFP and polyHIV (green) and dCasRx-GFP polyNT (orange) transfected cells. The means ± s.e.m. are depicted as horizontal and vertical bars for each group, respectively. Student’s t-test was used for statistical comparisons, and two-tailed *p* values are indicated.

**Supplementary Figure S5.**
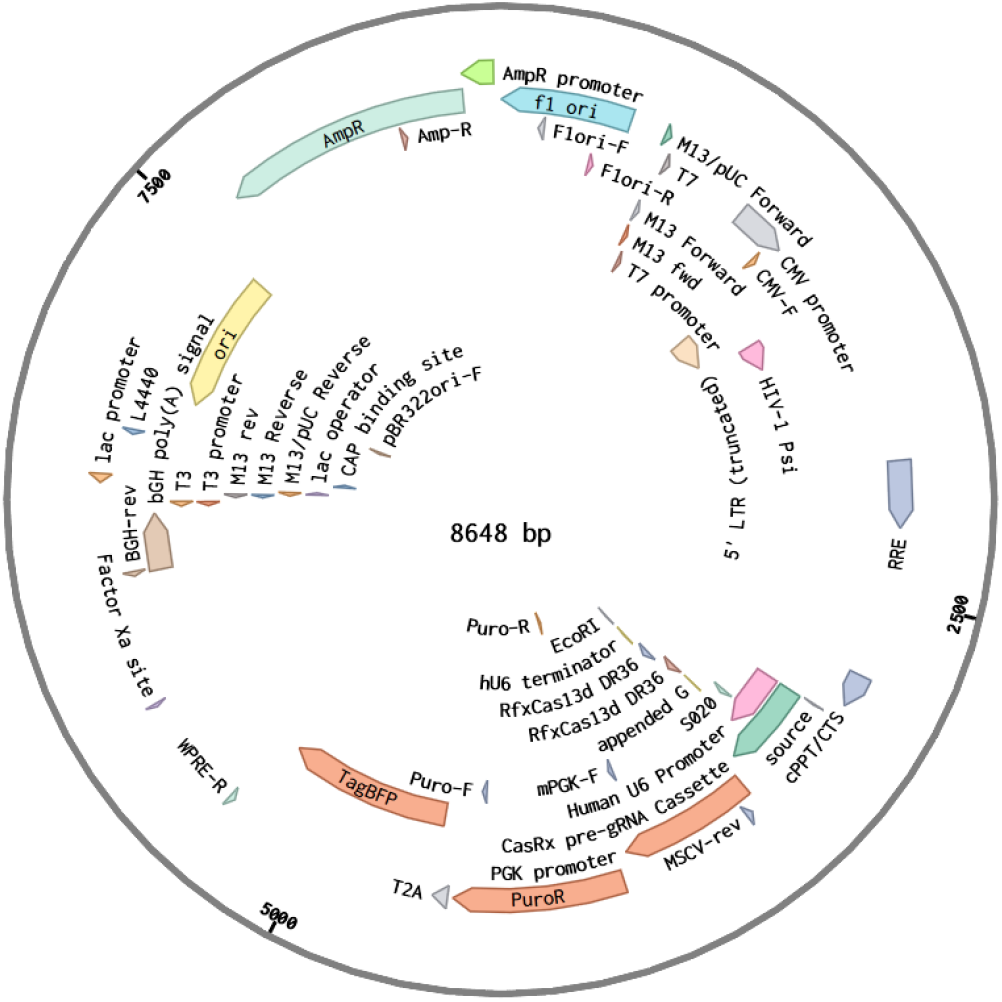
Plasmid Design – pKLV2-U6-CasRx-(pre-gRNA)-PGKpuro2ABFP-W.

